# Dynamical Biomarkers of Creative Cognition Across Divergent and Convergent Problem-Solving

**DOI:** 10.64898/2026.05.03.722474

**Authors:** Anubhav, Tzu-Ling Liu, Yang Li, Kazuyuki Aihara, Kantaro Fujiwara, Zenas C. Chao

## Abstract

Creativity fluctuates markedly from moment to moment, even within the same individual, yet the neural dynamics that determines whether a given attempt produces a highly creative idea remains poorly understood. Prior studies have identified static EEG correlates of creative thinking, but these do not explain how brain activity is dynamically organized before and during successful idea generation. Here, we model creative cognition as trajectories through a neural state space using energy landscape analysis (ELA) of EEG recorded during two complementary problem-solving paradigms: the divergent Alternative Uses Test (AUT) and the convergent, goal-directed Fusion Innovation Test (FIT). Across both tasks, creative success was associated with dissociable dynamical signatures in the resting and ideation stages. Before ideation, greater diversity of resting-state patterns of activity, corresponding to possible attractors, indexed a preparatory substrate of creative potential, showing weak trial-level effects but robust subject-level coupling with performance. During ideation, higher creativity was predicted by how the brain traversed its accessible state space: successful trials were characterized by traversal biased toward sustained exploration within stable attractor basins rather than frequent switching between basins of attraction (***β* = 0.104, *p* = 0.004**, trial-level). These findings identify a task-invariant, biologically grounded dynamical mechanism of creative cognition and show that creative performance depends not only on which neural states are available, but also on how neural activity traverses that state space.

## 1 Introduction

Creativity is often defined as the capacity to generate ideas that are both novel and useful [1, 2], yet the underlying cognitive and neural mechanisms remain difficult to pin down because creative performance is neither constant nor purely trait-like. Even within the same individual, creative output fluctuates across time and context, suggesting that moment-to-moment brain dynamics may shape the likelihood that a given attempt yields a creative solution [3, 4]. A mechanistic account therefore requires biomarkers that capture not only which neural features are present, but how neural activity organizes and evolves as the mind moves from a pre-task baseline into active idea generation [5].

Electroencephalography (EEG) is well suited to this problem because it provides a direct window onto fast neural dynamics [6]. Prior work has identified reproducible EEG correlates of creative thinking, most notably task-related synchronization in the alpha frequency band (8–12 Hz) [7–9], which is thought to reflect internally directed attention and the inhibition of task-irrelevant sensory information [10, 11]. However, much of the EEG creativity literature has focused on time-averaged power, connectivity, or univariate contrasts between task and baseline phases [12, 13]. Such static summaries risk obscuring the temporal structure of the underlying neural processes, even though creativity is thought to unfold as a sequence of complex transitions between cognitive states [14–16].

To address the limitations of time-averaged measures, recent studies have begun to adopt dynamic approaches, such as Hidden Markov Models (HMM) and microstate analysis, to capture the temporal evolution of brain states during creative tasks like metaphor generation and design ideation [4, 17, 18]. While these methods successfully identify sequences of neural states, including transitions associated with idea novelty, they do not explicitly quantify state stability or the geometry of switching between states. In this study, we address this gap by modeling creative cognition as trajectories on a neural state landscape and deriving interpretable dynamical biomarkers from EEG using energy landscape analysis (ELA) (Fig. 1). This probabilistic perspective aligns with recent theories viewing design creativity as a nonlinear, chaotic process sensitive to initial conditions [19], while ELA provides a principled bridge between the theoretical dynamics and the geometry of the underlying landscape [20–22].

**Fig. 1:**
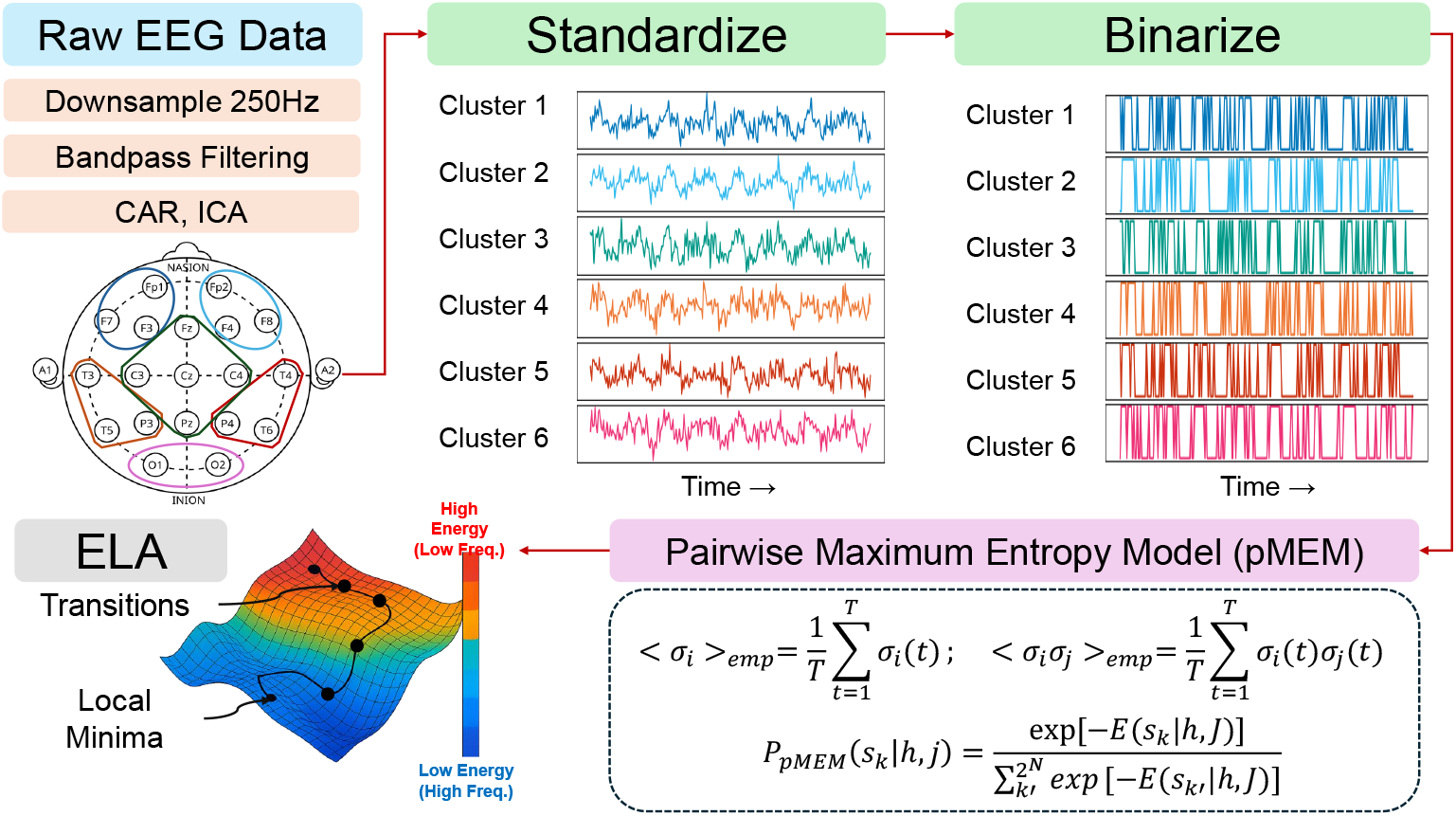
Energy landscape analysis pipeline. In this study, EEG preprocessing was followed by spatial clustering, within-trial standardization, and binarization to form a state sequence. Later, these binary state sequences were used to fit a pairwise maximum entropy model (pMEM), yielding an energy landscape from which local minima (attractor basins) and transition metrics were computed.

Thus, our primary hypotheses are stage-specific, with distinct predictions for resting-state and ideation-phase dynamics, and conceptualize creative cognition as trajectories in a neural state space. This is consistent with findings that intrinsic resting-state variability [23, 24] and preparatory neural activity [25] serve as possible markers of “creativity potential” [5], suggesting that greater pre-task attractor diversity may support subsequent creative output, particularly when the task demands generate nonlinear dynamics with divergent and convergent modes. In contrast, ideation-phase dynamics may reflect ongoing reconfiguration of large-scale networks (e.g., Default Mode Network and Executive Control Network) [16] and a richer exploration of state space once the task is underway.

While neural analyses must respect the dynamic structure of creative cognition, a fundamental challenge arises from the behavioral assays typically employed. The Alternative Uses Test (AUT) has long served as the gold standard for probing divergent thinking [26, 27]. However, this monolithic approach represents a significant methodological blind spot: neural dynamics observed in AUT-only studies may merely reflect the task-specific mechanics of unconstrained verbal generation and semantic retrieval, rather than the core mechanisms of creativity itself [28, 29]. Real-world creative problem solving rarely occurs in a vacuum of unconstrained generation; it necessitates convergent evaluation to narrow the search space and select effective solutions [30]. Indeed, recent theoretical frameworks suggest that creative cognition is best understood not as a dichotomy but as a dynamic continuum that fluctuates between divergent and convergent modes [31]. To overcome the limitations of the single-task paradigm, the present study employs a dual-task design, pairing the unconstrained AUT with the recently introduced Fusion Innovation Test (FIT). The FIT requires participants to combine distinct elements to achieve a specific goal, thereby engaging both the generative and evaluative components of creativity [32].

Comparing neural signatures across these two tasks allows us to isolate markers that generalize across distinct creative demands, ensuring that they reflect task-independent creative cognition rather than the idiosyncrasies of a single paradigm. Figure 2 illustrates the task logic and scoring dimensions, where novelty and feasibility are shared by both tasks, but there is an additional goal criterion in FIT [32, 33]. In parallel, the integration of automated scoring approaches has begun to resolve longstanding limitations in creativity measurement, such as rater subjectivity and the inability to process large datasets [34]. Accordingly, we use task ratings from large language models (LLMs), which have been shown to reliably quantify multiple dimensions of creative quality and to perform comparably to, or in some cases better than, human raters [33].

**Fig. 2:**
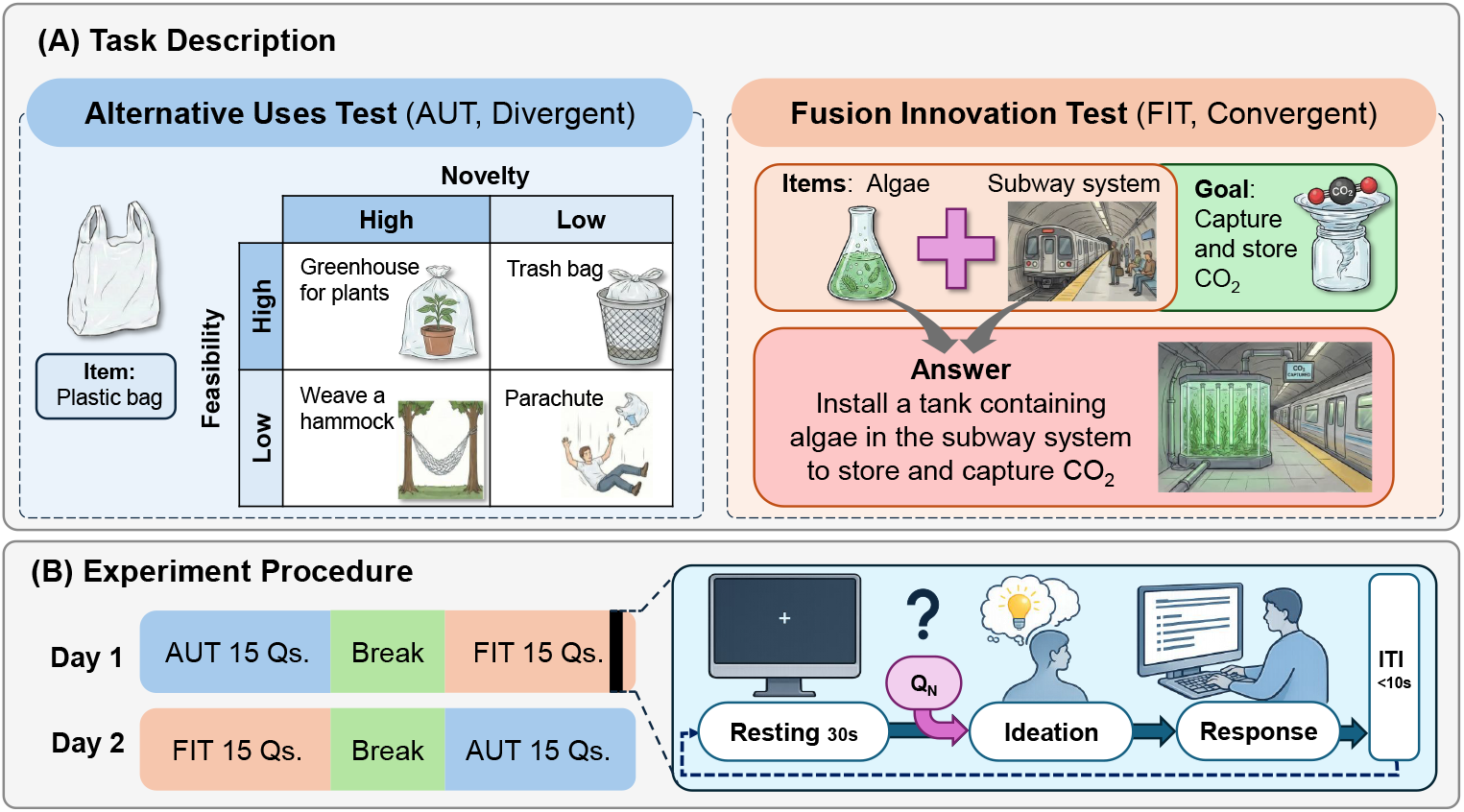
Task Description and Experiment Procedure. AUT prompts participants to generate alternative uses for a common object, emphasizing novelty and feasibility. FIT prompts participants to fuse two items to achieve a stated goal, emphasizing novelty, feasibility, and goal effectiveness. Across days, AUT and FIT blocks were counterbalanced. Within each trial, a 30 s eyes-open resting segment directly preceded the ideation phase; the response phase occurred after ideation and was not included in the analyzed EEG.

Overall, our work makes three contributions. First, it introduces a dynamical, state-space account of EEG markers of creativity that complements predictive feature-based approaches by linking creative performance to attractor topology and transition dynamics. Second, it explicitly distinguishes pre-task resting-phase dynamics from ideation-phase dynamics within the same trials, allowing us to test a staged mechanism in which preparatory attractor diversity and ideationasl traversal play dissociable roles. Third, we demonstrate that, even with the complementary demands of AUT and FIT tasks, the dynamical biomarkers are task-invariant and govern creative success across the entire divergent-convergent continuum.

## 2 Results

### 2.1 H1: Resting-phase attractor diversity shows a weak association with creative performance

We first tested whether resting-phase landscape topology and traversal relate to trialwise creativity (H1). Descriptively, creativity (question-normalized *z*-score) increased with resting-phase LMC across pooled trials (Spearman *ρ* = 0.08, *p* = 0.001, *N* = 1647 trials), with a stronger trend in FIT than AUT (Fig. 3 (a)). In contrast, the number of resting-phase transitions showed near-zero association with creativity (Fig. 3 (b)). These descriptive patterns suggest that larger resting attrasctor repertoires may relate to higher performance, whereas rapid resting switching is not informative at the trial level.

**Fig. 3:**
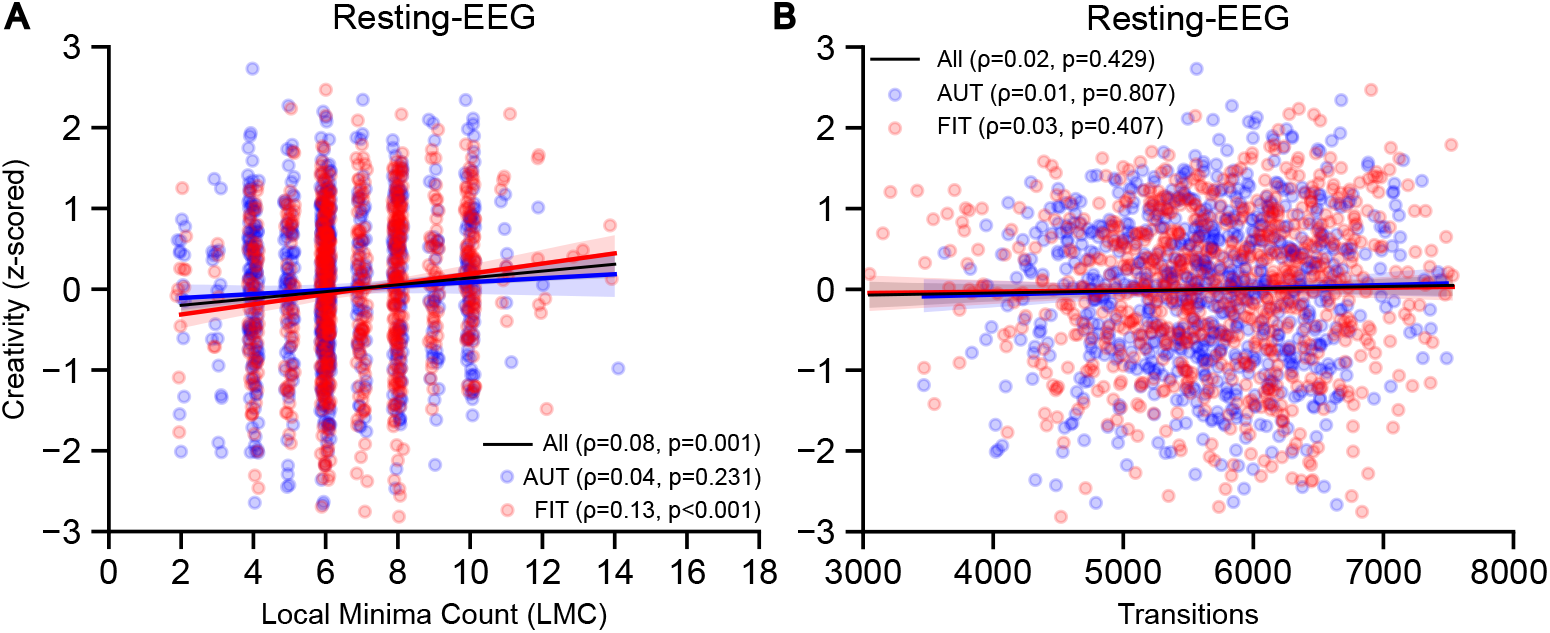
Resting-phase associations between landscape metrics and creativity. Scatter plots show trial-wise creativity (question-normalized *z*-score) versus (a) the local minima count (LMC) and (b) the number of transitions during the 30 s resting segment. Lines indicate the task-specific trends (AUT, FIT) and pooled trend (All).

Primary inference accounted for repeated measures using mixed-effects models with random intercepts for subjects and questions. For resting-phase LMC, the following trial-level mixed model was investigated:

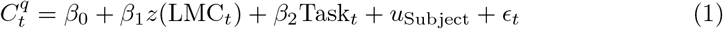

yielding a weak, non-significant fixed effect of LMC (*β*_1_ = 0.020, *p* = 0.445; 95% CI [−0.032, 0.073]). Since the trial-level association between resting LMC and creativity was not supported after accounting for nesting, we suspect that resting-phase topology metrics are largely subject-specific.

### 2.2 H2: Ideation-phase traversal mode predicts trial-wise creativity

We next tested whether ideation-phase landscape topology and traversal relate to creativity (H2). Descriptively, while ideation-phase LMC showed little relationship with creativity, transitions exhibited a positive association, with stronger trends in FIT than AUT (Fig. 4). However, transition counts covary strongly (e.g., *r* = 0.97 between transitions and EEG length), and after accounting for duration adjustments, neither transitions nor length-normalized transition rates showed a reliable trial-wise association with creativity. This motivated a shift from *how much* traversal occurred to *what kind* of traversal dominated during ideation.

**Fig. 4:**
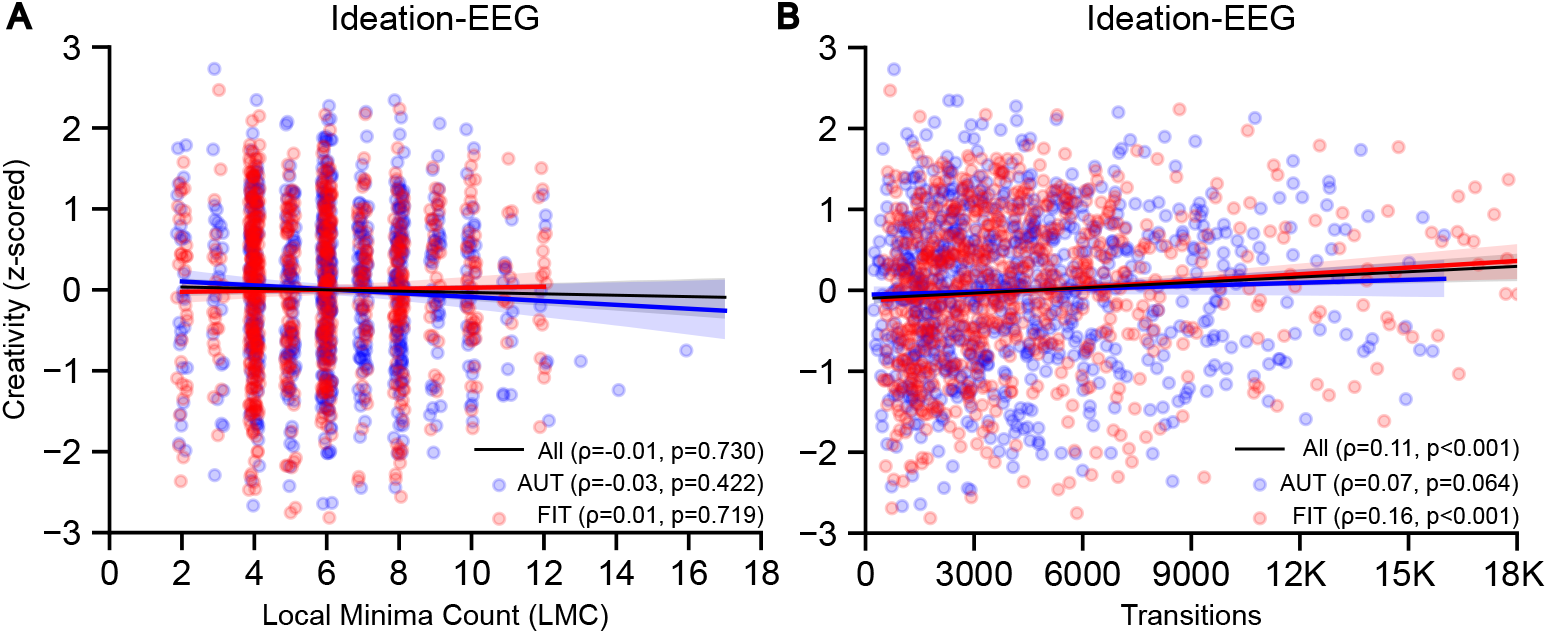
Ideation-phase associations between landscape metrics and creativity. Scatter plots show trial-wise creativity (question-normalized *z*-score) versus (a) ideation-phase LMC and (b) ideation-phase transitions. Lines indicate the task-specific trends (AUT, FIT) and pooled trend (All).

We therefore quantified a traversal mode as the balance between within-basin and between-basin transitions using a within-basin traversal bias (WBTB) metric as follows:

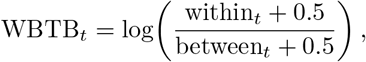

and modeled creativity (question-normalized z-score) using a linear mixed-effects model with a subject-level random intercept and fixed effects of traversal bias and Task. Because within-basin traversal bias was effectively uncorrelated with ideation EEG length (*r* = *−*0.03), we did not include EEG length as a covariate in this model:

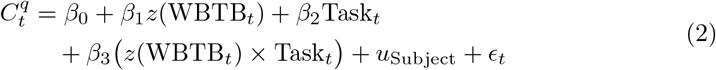

Within-basin traversal bias robustly predicted creativity across both tasks (*β*_1_ = 0.104, *p* = 0.004), with trials exhibiting greater within-basin traversal (and fewer between-basin transitions) consistently yielding higher question-normalized creativity scores. Crucially, the predictive power of the within-basin traversal bias was highly generalizable. The Task main effect was negligible (*β*_2_ = −0.004, *p* = 0.929), and the mixed-effects model revealed no significant interaction with Task (*β* = −0.065, *p* = 0.182), indicating that the biomarker’s predictive power held equally across both paradigms. These results refine the traversal-based account of creative ideation: rather than being driven by faster switching per se, higher-quality ideas were associated with a traversal mode biased toward sustained within-basin dynamics during ideation, regardless of whether the task demanded divergent thinking (AUT) or convergent evaluation (FIT).

### 2.3 Stage-dissociable dynamical biomarkers of creativity

The results are consistent with a stage-dissociable account of creative performance in this dataset. At rest (H1), attractor diversity (LMC) operates as a stable, between-subject marker. While rigorous block bootstrapping confirmed that its trial-level association with creativity is not significant after accounting for nested variance (*β* = 0.009, 95% CI [-0.053, 0.068], bootstrap *p* = 0.746), a subject’s overall baseline LMC reliably predicts their mean creative performance across all tasks (*β* = 0.411, *p* = 0.030). In contrast, during active ideation (H2), creative performance robustly tracked state-level shifts in the *traversal mode*: individual trials characterized by relatively greater within-basin dynamics yielded significantly higher creativity scores (*β* = 0.119, 95% CI [0.032, 0.212], bootstrap *p* = 0.006).

Importantly, resting topology and ideation dynamics were coupled at the subject level: higher resting LMC predicted a shift in the average ideation traversal mode (lower within-basin traversal bias) (*β* = −0.575, *p <* 0.001, task interaction *ns*). These findings suggest separable contributions of (i) a trait-like resting substrate indexed by attractor diversity and (ii) a state-like ideation process indexed by traversal mode. Figure 5 summarizes this staged interpretation: lower mean creativity across individuals was associated with lower resting LMC, whereas within individuals, higher trial-wise creativity during ideation was associated with more within-basin-biased traversal.

**Fig. 5:**
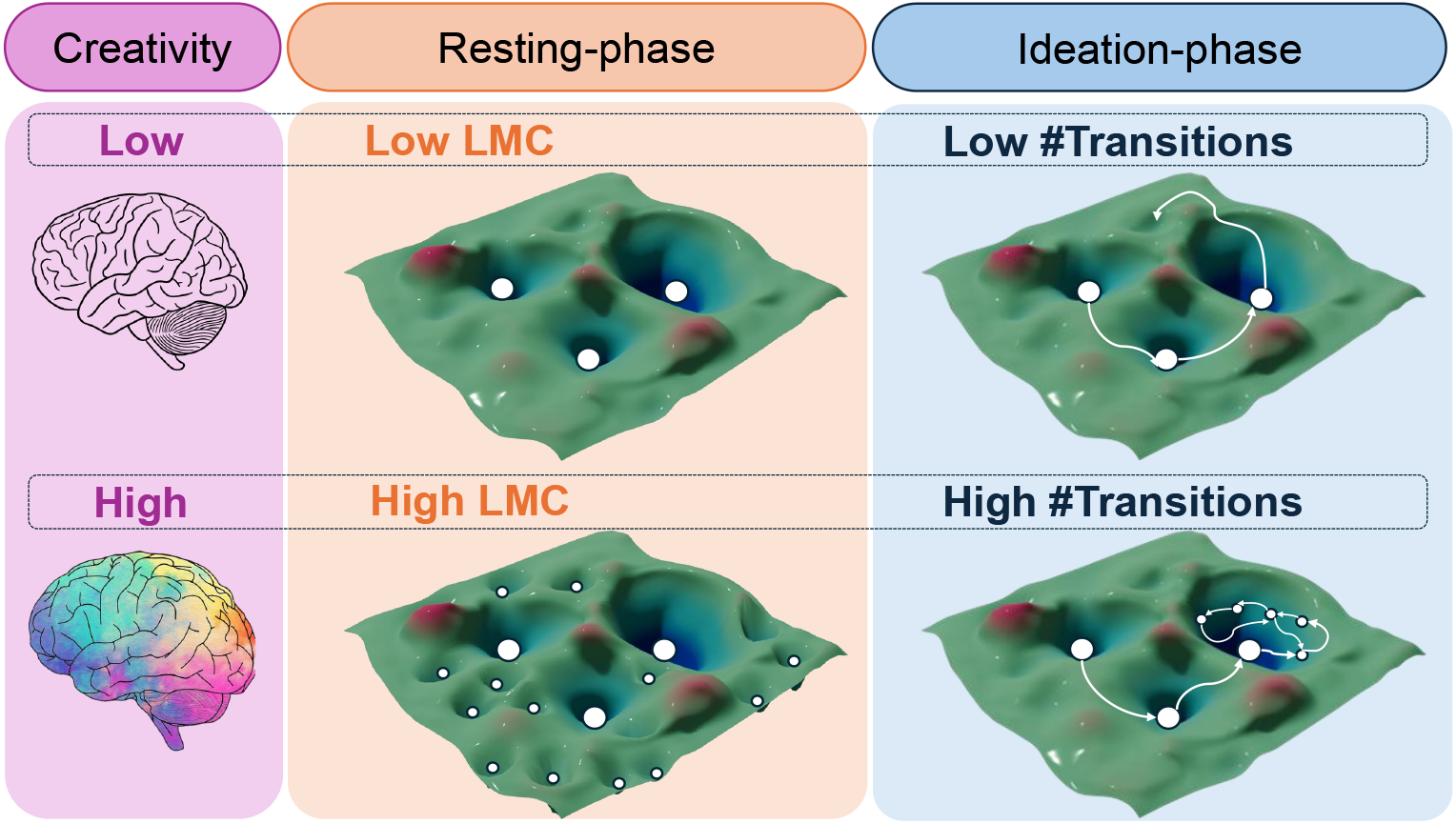
Conceptual summary of staged dynamical biomarkers. Schematic illustration of the hypothesized two-stage interpretation: resting-phase attractor diversity (LMC) reflects the availability of dynamical options prior to ideation, whereas ideation-phase within-basin traversal bias reflects traversal dynamics during active idea generation.

## 3 Discussion

This study applies an energy landscape framework to trial-resolved EEG to test whether dynamical organization of brain states predicts creative performance across two distinct problem-solving tasks (AUT and FIT) and two stages within each trial (resting and ideation). Results are consistent with a stage-based account in which resting attractor diversity reflects a preparatory substrate that varies primarily across individuals, whereas ideation success is most tightly coupled to how the system traverses its accessible state space during active idea generation.

### 3.1 Resting-phase attractor diversity as a weak but principled preparatory signature

The clearest resting-phase signal emerged at the between-subject level (*β* = 0.141, *p* = 0.038). This aligns with prior research indicating that individual differences in resting-state functional connectivity and temporal variability are significant predictors of creative potential [35]. It is also compatible with the relatively short resting segment (30 s) and with the possibility that resting dynamics relates to creative performance by shaping how efficiently participants enter a productive ideation regime rather than by directly predicting idea quality. In line with this suggestion, we observed a clear cross-stage coupling at the subject level: resting LMC predicted the average ideation traversal mode (task-invariant, *β* = −0.578, *p <* 0.001), indicating that resting topology covaries with subsequent ideation traversal style at the subject level. This stage distinction is consistent with the notion that preparatory state properties bias subsequent processing strategies (e.g., insight vs. analysis) [36], while ideation-phase dynamics more directly reflects the generative process itself [25].

### 3.2 Ideation-phase traversal as a stage-specific dynamical marker of creativity

Ideation performance was more closely related to the *traversal mode* than to overall switching frequency. In the energy landscape view, this pattern suggests that higher-quality ideation is linked less to faster state switching per se and more to a traversal regime biased toward sustained dynamics within basins. This complements prior work showing that preparatory EEG signatures predict trial-to-trial creative performance across AUT and FIT [5] by specifying an interpretable dynamical property expressed during the ideation window. One possibility is that within-basin-biased traversal reflects internally directed processing states that support controlled retrieval and integration, consistent with prior links between creative cognition and oscillatory signatures of internal attention [10]; testing this correspondence will require further analyses that explicitly relate the traversal mode to frequency-resolved or network-level markers.

### 3.3 A task-invariant mechanism of creative cognition

A long-standing controversy in the cognitive neuroscience of creativity concerns whether creative cognition relies on domain-general or task-specific neural mechanisms [37–40]. Because brain dynamics are highly sensitive to task demands, it has been difficult to determine whether observed neural signatures reflect the fundamental process of creative thought or merely the cognitive requirements of task compliance [41, 42]. Our finding that a within-basin-biased traversal mode predicts trial-wise creativity across both the AUT and the FIT offers a principled resolution. Since these tasks impose fundamentally different cognitive demands (unconstrained divergent generation versus goal-directed convergent integration), a traversal signature that survives across both is unlikely to be an artifact of task-specific compliance.

This points to a task-invariant, domain-general mechanism of creative thought grounded in the dynamical traversal of neural state space. The ideation-phase signature identified here complements the shared preparatory marker of creativity potential reported across the same tasks [5]; together, they support a continuous rather than dichotomous view of divergent and convergent thinking [31], suggesting that the core substrate of creative ideation is a task-independent capacity to sustain exploration within stable neural attractors.

### 3.4 Limitations and future directions

Several limitations bound the interpretation and suggest clear next steps. First, state transitions were computed from the sign of the preprocessed EEG signal. Since in ELA, binarization was applied after band-pass filtering, common average referencing, and ICA-based artifact correction, the resulting trajectories are likely less affected by artifactual zero-crossing jitter than unprocessed voltage traces. Even so, sign-based binarization imposes a coarse representation of neural activity and may remain sensitive to analytic choices. Alternative state-space methods, including HMM and microstate approaches, may capture more temporally sustained structure and thus provide complementary estimates of neural dynamics [4, 43]. Future work should test whether the present traversal metrics are robust to temporal coarse-graining and to variation in the binarization threshold, and determine how these choices affect inferred landscape geometry and transitions.

Second, ideation duration varied substantially across tasks, reflecting differences in allotted task time (AUT: 90 s; FIT: 240 s). Accordingly, for the ideation phase, we adjusted EEG segment length in the trial-level models and applied a log-ratio traversal metric to mitigate simple time-on-task effects; however, these duration differences may still influence the trial-wise pMEM landscape estimation. Future work can address this by analyzing duration-matched windows, estimating traversal metrics in sliding windows to test whether the traversal mode evolves over the ideation period or designing protocols with matched ideation durations across tasks.

Third, the energy landscapes were estimated at the sensor level after spatial clustering into six regions, a coarse-grained representation that trades anatomical specificity for statistical tractability at the single-trial level. Importantly, however, the dominant local minima identified by the pMEM are not arbitrary computational constructs. A post hoc topographic validation confirmed that they organize into biologically meaningful families with strong correspondence to canonical *D*-like, *B*-like, and *C*-like microstate patterns (Appendix A.2), lending direct neurophysiological support to the energy landscape representation. Rather than viewing the absence of a strict one-to-one microstate mapping as a limitation, we suggest this reflects a conceptual advance: whereas canonical microstates are derived purely by geometric clustering of scalp topographies [44], the pMEM local minima are grounded in the dynamical structure of the energy landscape, identifying recurrent brain states by virtue of their attractor stability rather than their spatial similarity alone. This dynamically principled state representation offers a novel perspective on resting-state EEG organization, and future work could systematically characterize how pMEM attractors relate to the full microstate taxonomy across cognitive contexts. The sensor-level formulation does, nonetheless, limit the anatomical precision with which attractors can be mapped to cortical systems; higher-density EEG, source reconstruction, or multimodal approaches such as EEG–fMRI [19] could further test how specific cortical networks generate attractor topology and traversal dynamics [45].

Despite these limitations, the present results demonstrate that energy-landscape-derived dynamical biomarkers can link trial-wise EEG dynamics to creativity across divergent and convergent tasks. The strongest evidence supports a stage-specific mechanism in which ideation-phase traversal dynamics index as creative success (H2), while resting-phase attractor diversity provides a weaker preparatory signature (H1). These findings provide a foundation for future work that combines dynamical state-space modeling with richer behavioral decomposition of ideation and evaluation processes. Ultimately, such metrics could inform the development of real-time neurofeedback systems capable of detecting when designers transition between exploratory and evaluative states, potentially aiding in the modulation of creative performance.

## 4 Methods

### 4.1 Dataset Description

Twenty-eight healthy participants (17 males, 11 females; mean age 22 ± 2.5 years, range 18–29) each completed two creativity tasks: the Alternative Uses Test (AUT) and the Fusion Innovation Test (FIT) (Fig. 2). This yielded 1,647 analyzable trials 10 in total (AUT: 813; FIT: 834), with each participant contributing between 48 and 60 trials. Each trial unfolded in three successive segments: a 30 s eyes-open resting period, an ideation period during which participants silently generated responses, and a typed response period. EEG analyses were restricted to the resting and ideation periods; the response period was excluded to avoid motor-related confounds.

### 4.2 Creativity scoring

Creativity was assessed at the trial level using an LLM-based automated scoring pipeline previously validated for AUT and FIT responses [32, 33]. Each response was rated along multiple dimensions: `Novelty` and `Feasibility` for AUT trials, and `Novelty, Feasibility`, and `Goal` for FIT trials. Trial-level creativity was computed as the geometric mean of the available dimensions:

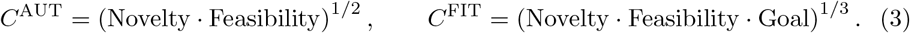

To normalize across prompt difficulty and scale differences, we computed a question- normalized creativity score within each task:

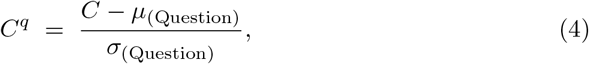

where *µ*_(Question)_ and *σ*_(Question)_ denote the mean and standard deviation computed across all trials for the same `Question`.

### 4.3 Data Preprocessing

EEG was recorded from 19 channels arranged according to the 10–20 system. Signals were preprocessed using a standard pipeline (Fig. 1): downsampled to 250 Hz, banpass filtered between 0.5 and 100 Hz, re-referenced to the common average, and cleaned via independent component analysis (ICA) to remove artifacts.

To enable trial-wise energy landscape estimation over a tractable state space, the 19 scalp electrodes were grouped into six spatial clusters, yielding a six-node binary representation with 2^6^ = 64 possible system states. This coarse-grained resolution was chosen to preserve large-scale scalp-field organization while keeping the pMEM tractable at the single-trial level. The clustering was specifically designed to capture broad anterior–posterior and lateralized voltage gradients, which underlie the dominant spatial patterns of canonical resting-state EEG microstates [44, 46]. Grouping the central-midline electrodes into a shared cluster additionally reduces sensitivity to zero-crossing jitter around the midline, so that binary state transitions more faithfully reflect global field reconfigurations rather than isolated channel fluctuations [47]. The neurophysiological plausibility of this representation is further evaluated through a post hoc topographic comparison of the dominant local minima against canonical EEG microstates, reported in Appendix A.2.

Let *x*_*c*_(*t*) denote the cluster-averaged voltage at time *t* for cluster *c* ∈ {1, …, 6}. Clusters were defined as follows:

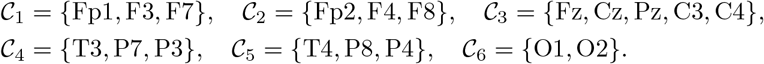

For each trial and each stage (rest; ideation), we z-scored each cluster time series across time as follows:

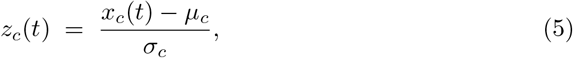

where *µ*_*c*_ and *σ*_*c*_ are computed within the given trial and stage.

We discretized the multivariate EEG into a binary state sequence by thresholding the standardized cluster signal at zero:

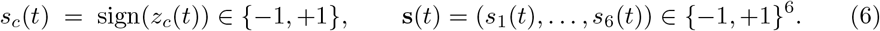

Thus, each time point maps to one of 2^6^ = 64 possible binary states. We retained the original 250 Hz temporal resolution for state transitions.

### 4.4 Pairwise maximum entropy model (pMEM)

For each trial and stage, we fitted a pairwise maximum entropy model to the observed binary state sequence 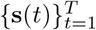, yielding an Ising-form energy function:

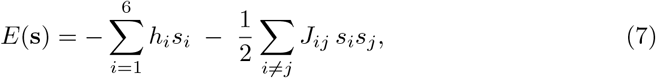

with corresponding probability

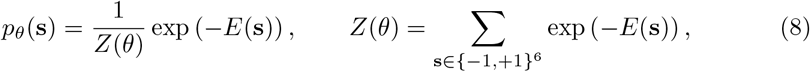

where *θ* = {*h*_*i*_, *J*_*ij*_ } are parameters.

Parameters were estimated via pseudo-likelihood maximization using gradient descent, with a maximum of 1000 iterations per fit. The pseudo-likelihood is given as:

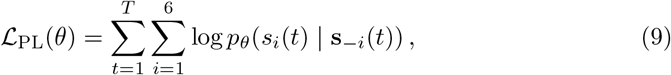

where **s**_*−i*_ denotes all spins except *i*. Model goodness-of-fit was additionally quantified using KL-divergence and entropy differences between empirical and model distributions.

#### Algorithm 1

Basin assignment by steepest descent

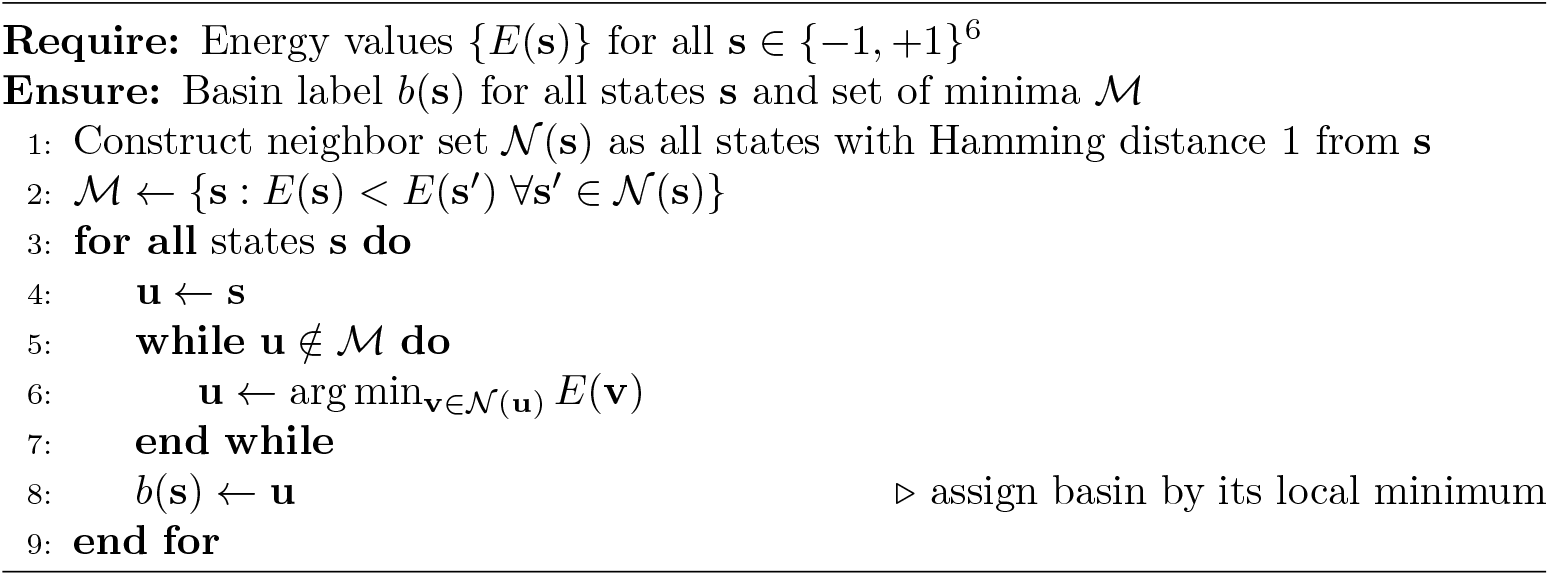

### 4.5 Energy Landscape Construction and Basin Assignment

Local minima were identified across all 64 states of *E*(**s**) using single-bit flip neighborhoods: a state **s** qualifies as a local minimum if its energy is lower than that of all Hamming-distance-1 neighbors (Algorithm 1). Each state was then assigned to an attractor basin via deterministic steepest descent, in which the system iteratively transitions to its lowest-energy neighbor until a local minimum is reached, yielding a unique basin label for every state. The local minima count (LMC), defined as |*ℳ*|, was extracted for each trial and stage. Representative examples of frequently visited states are illustrated in Fig. 6.

**Fig. 6:**
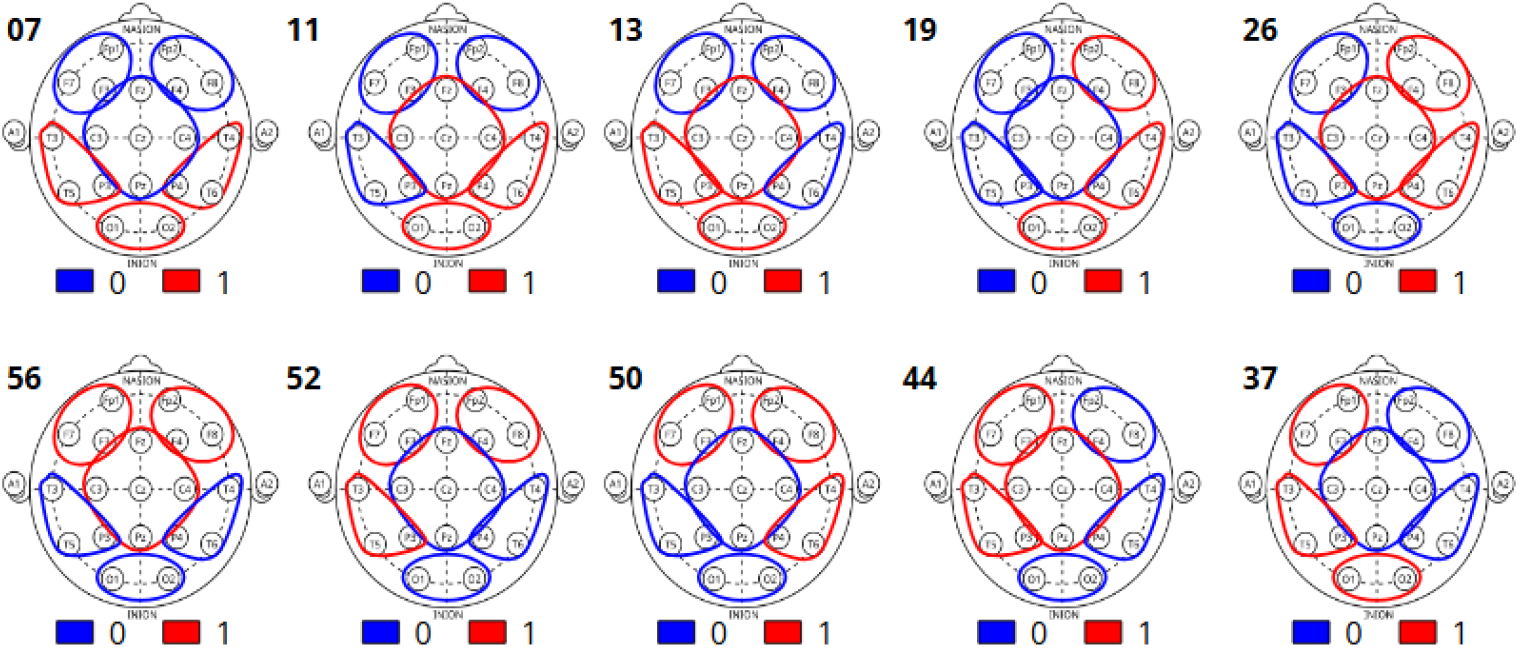
Representation of dominant local minima. Scalp maps visualize top ten most frequently observed local minimum states (with state IDs), illustrating structured patterns across the six spatial clusters used for ELA.

### 4.6 Traversal metrics

Let **s**(*t*) be the observed state sequence and *b*(**s**(*t*)) its basin label. We computed:

- **State transitions**

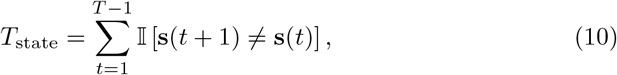 and for ideation, the transition rate *R*_state_ = *T*_state_*/*Δ*t* (transitions per second), where Δ*t* is stage duration.
- **Within-basin transitions**

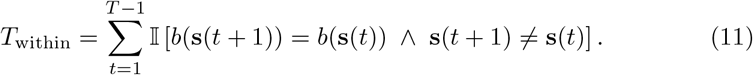
- **Between-basin transitions**

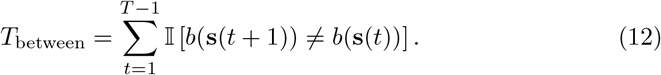

## Acknowledgements

This research is supported by JSPS KAKENHI grants JP22K18419, JP24K15161, and JP25H00451 (to K.F.), JST Moonshot RD grant JP-MJMS2021 (to Y.L., K.A., and K.F.); Beyond AI Joint Project, Institute of AI and Beyond of UTokyo (to Y.L. and K.A.); a project, JPNP14004, commissioned by the New Energy and Industrial Technology Development Organization (NEDO) (to K.A. and K.F.); World Premier International Research Center Initiative (WPI), MEXT, Japan (to Z.C.C.), IRCN-Daikin SCP (to Z.C.C.)

## Declarations

- Conflicts of interest: The authors declare that no conflicts of interest exist.
- Ethics approval and consent to participate: All participants provided their informed consent, and the study was approved by the research ethics committee.
- Code availability: The custom Python code used for all statistical analyses is publicly available on GitHub (https://github.com/anubhav2901/EEG-ELA-Creativity.git). The raw EEG data generated in this study are not publicly available due to ethical restrictions regarding participant privacy; however, a derivative, de-identified dataset is available from the corresponding author upon reasonable request.
- Author contribution: Z.C.C. and T.L.L. conceptualized and designed the research. T.L.L. performed the experiments. T.L.L. and A. analyzed the data. A. developed the methodology and prepared the figures. A., T.L.L., Y.L., and Z.C.C. contributed to data interpretation and validation. Z.C.C. supervised the project and acquired funding. A. wrote the original draft. All authors reviewed, edited and approved the final manuscript.

## Appendix A Supplementary Analyses

This section reports robustness analyses and mechanistic decompositions that complement the primary results. Unless noted otherwise, we use hierarchical inference (mixed-effects or fixed-effects with subject-block bootstrap as appropriate) and question-normalized creativity (*C*^*q*^; Eq. 4).

### A.1 Subject-level robustness (aggregation with FDR)

To provide a participant-level robustness check that avoids trial-level nesting concerns, we aggregated each ELA metric and creativity score by participant (mean across trials) within stage, then computed Spearman correlations between subject-level averages and creativity. *p*-values were corrected using Benjamini–Hochberg FDR across the tested features within each stage.

At the subject level, resting-phase LMC showed a positive trend with creativity (*ρ* = 0.355, *p* = 0.064, *q* = 0.127), whereas resting transitions were not associated (*ρ* = −0.088, *p* = 0.656; Table A1). During ideation, transitions showed a positive trend (*ρ* = 0.401, *p* = 0.035, *q* = 0.069), consistent in direction with descriptive trial-level patterns, but sensitive to trial duration, while ideation LMC showed a weaker association (Table A1). These subject-level analyses are conservative and reduce power; accordingly, they are presented as robustness checks rather than primary inference.

**Table A1:** Subject-level robustness correlations (Spearman) with FDR correction. Subject-level metrics were computed as the mean across trials per participant within stage; *q* denotes Benjamini–Hochberg FDR-corrected *p*-values within each stage.

| Stage | Feature | $\rho$ | $p$ | $q$ |
| --- | --- | --- | --- | --- |
| Resting | LMC | 0.355 | 0.064 | 0.127 |
| Resting | Transitions | -0.088 | 0.656 | 0.656 |
| Ideation | LMC | 0.201 | 0.304 | 0.304 |
| Ideation | Transitions | 0.401 | 0.035 | 0.069 |

### A.2 Topographic validation of the dominant local minima

To assess the neurophysiological plausibility of the dominant local minima identified by the pMEM, we compared their scalp topographies against the four canonical EEG microstates (*A*–*D*) [44, 46]. Although the energy landscape was estimated using a coarse-grained six-cluster state representation, the most frequently occupied local minima nonetheless exhibit large-scale topographies consistent with those of canonical resting-state microstates.

The ten most frequently occupied local minima in the resting-phase data (state IDs: 56, 7, 50, 13, 11, 52, 19, 44, 26, and 37, in descending order of occupancy) were selected for topographic analysis. As the canonical microstate reference, we used `Koenig 2002.set` from the Microstate Template Editor and Explorer repository^1^. For each participant and local minimum, time points whose binarized activity pattern in the dimension-reduced six-channel space matched that local minimum state were mapped back to the original preprocessed 19-channel EEG, and global field power (GFP)-peak scalp maps from those time points were extracted and averaged to yield a subject-level mean topography. Global-averaging across participants revealed five polarity-invariant families, each comprising a sign-reversed pair: (7, 56), (13, 50), (11, 52), (19, 44), and (26, 37) (Fig. A1). At the group level, these families exhibited the strongest spatial correspondence with microstate *D* for (7, 56), microstate *B* for (13, 50) and (11, 52), and microstate *C* for (19, 44), while (26, 37) showed only weak correspondence to any canonical template (Fig. A2a).

**Fig. A1:**
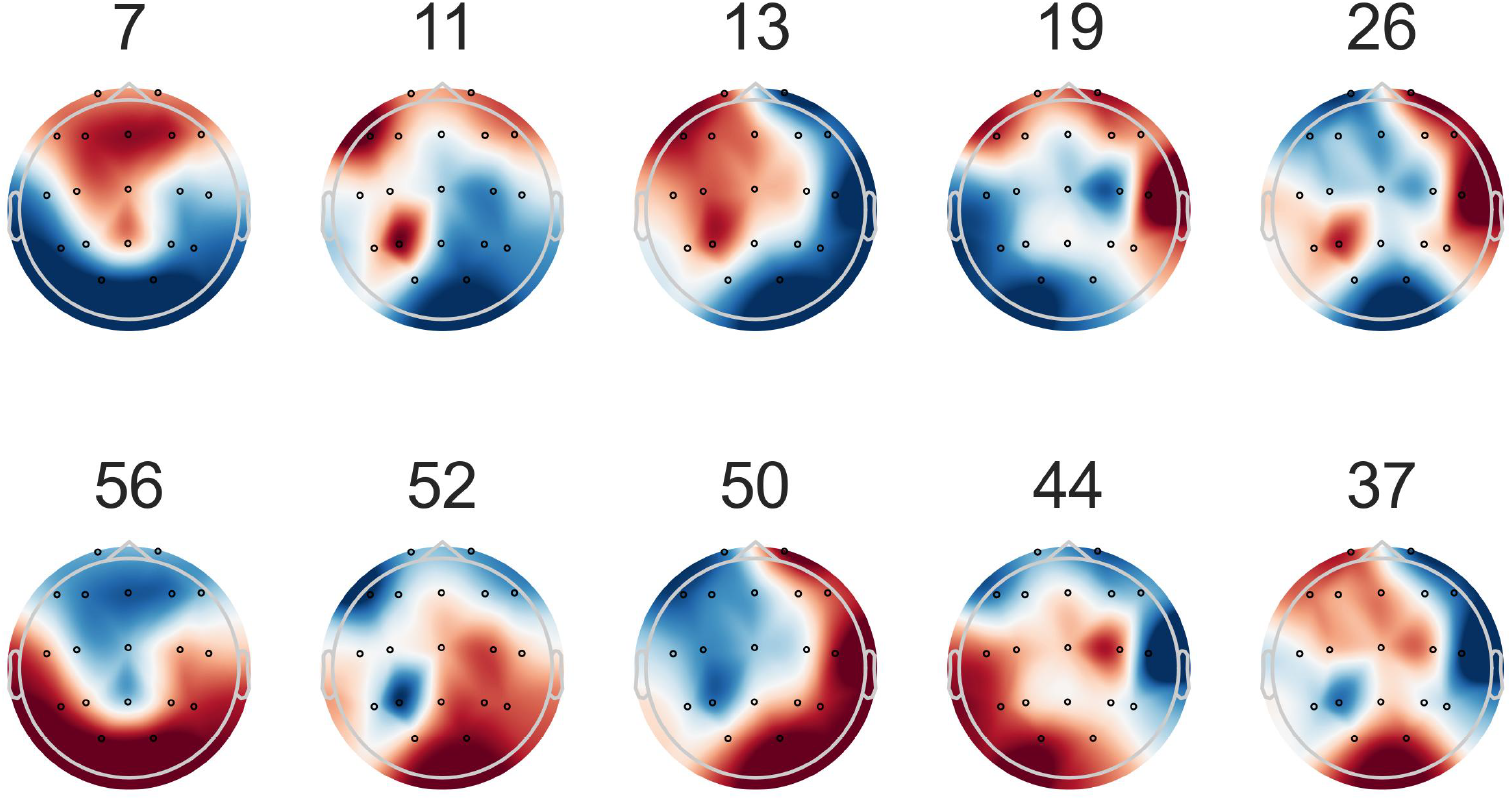
Global-average scalp topographies of the ten dominant local minima. The dominant local minima form five polarity-invariant families, each represented by a sign-reversed pair of maps: (7, 56), (11, 52), (13, 50), (19, 44), and (26, 37). These paired inversions support the interpretation that the ten dominant local minima reduce to five underlying topographic families.

**Fig. A2:**
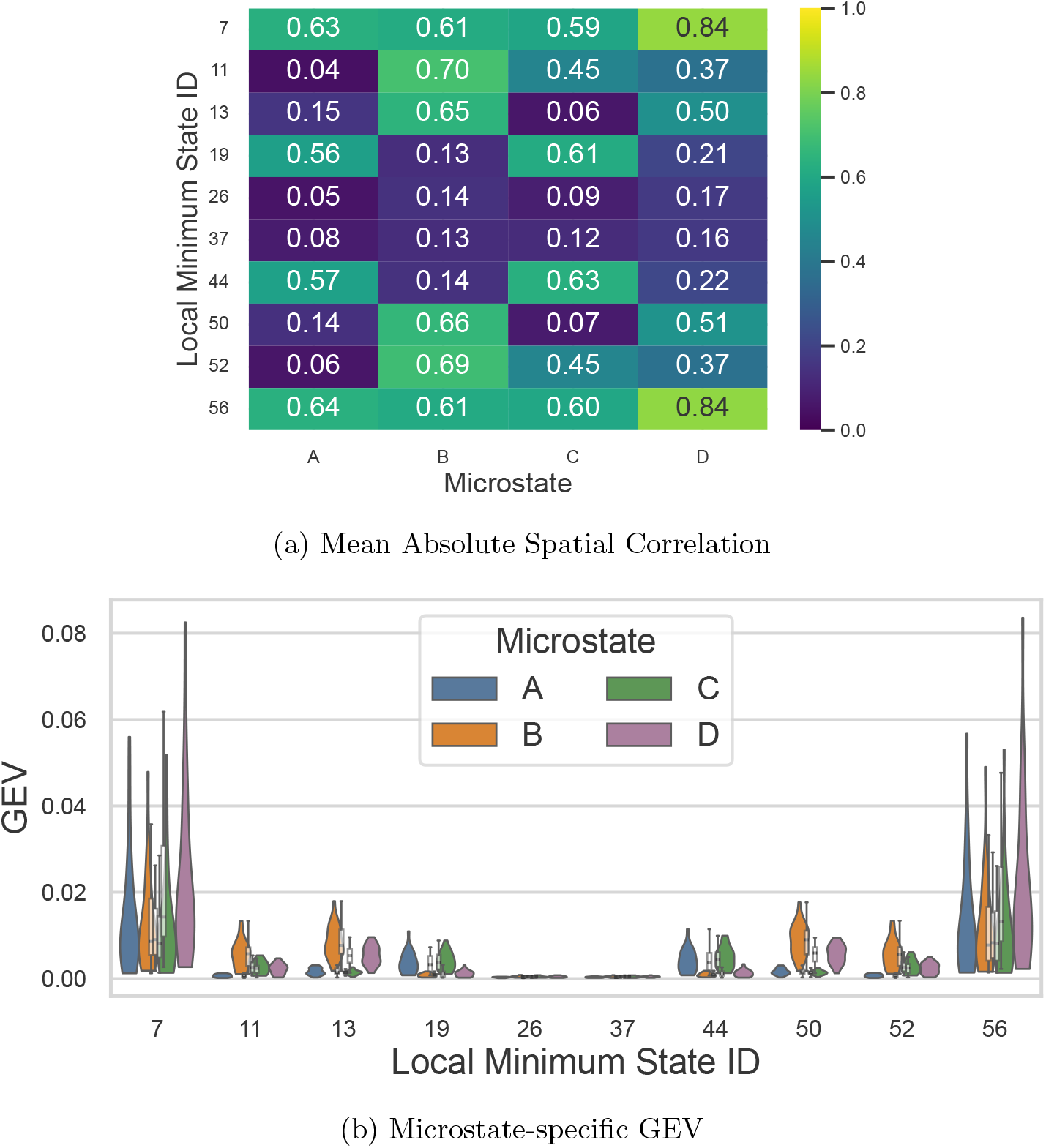
Topographic correspondence between dominant local minima and EEG microstates. (a) Group-level mean polarity-invariant spatial correlation between each dominant local minimum and the four microstates (*A–D*). (b) Microstate-specific GEV distributions across subjects for each l ocal minimum, showing the proportion of GFP-weighted variance explained by each microstate within the EEG samples assigned to that state.

Similarity to each canonical microstate template was quantified using polarityinvariant spatial correlation,

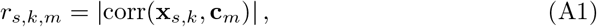

where **x**_*s,k*_ is the subject-level scalp map for subject *s* and local minimum *k*, and **c**_*m*_ is microstate template, where *m* ∈ {*A, B, C, D*}. We used the absolute value because EEG scalp fields are defined up to polarity reversal.

For each local minimum *k*, we tested whether the best-matching microstate was more strongly expressed than the second-best microstate across subjects. Let

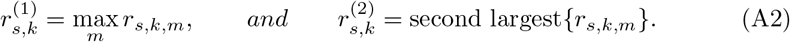

We then tested the directional contrast

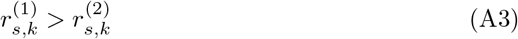

using one-sided Wilcoxon signed-rank tests across subjects, followed by Benjamini– Hochberg FDR correction across the ten local minima.

All ten dominant local minima exhibited significant median best-versus-second-best microstate separation after FDR correction (Table A2), though the magnitude of separation varied substantially across families. The clearest separation was observed for the *B*-like family (11, 52), followed by the *D*-like family (7, 56). By contrast, the *C*-like family (19, 44) displayed weaker separation, indicating greater microstate ambiguity, while the low-prevalence family (26, 37) yielded only small effect sizes despite reaching significance.

**Table A2:** Primary topographic validation: subject-level separation of median best- and second-best microstate matches. For each local minimum, subject-level scalp maps were matched to canonical EEG microstates (*A–D*) using polarity-invariant spatial correlation. The primary test compared the median bestversus second-best-matching microstate across subjects using a one-sided Wilcoxon signed-rank test. *q* denotes the Benjamini–Hochberg FDR-corrected *p*-value across the 10 local minima.

| State ID | $N$ | Median best | Median second | Median diff. | $p$ | $q$ |
| --- | --- | --- | --- | --- | --- | --- |
| 7 | 27 | 0.851 | 0.656 | 0.196 | $7.45e-9$ | $9.31e-9$ |
| 11 | 28 | 0.719 | 0.458 | 0.247 | $3.73e-9$ | $7.45e-9$ |
| 13 | 28 | 0.657 | 0.497 | 0.151 | $3.73e-9$ | $7.45e-9$ |
| 19 | 27 | 0.619 | 0.566 | 0.055 | $7.45e-9$ | $9.31e-9$ |
| 26 | 13 | 0.192 | 0.150 | 0.058 | $1.22e-4$ | $1.22e-4$ |
| 37 | 15 | 0.195 | 0.153 | 0.046 | $3.05e-5$ | $3.39e-5$ |
| 44 | 27 | 0.632 | 0.574 | 0.060 | $7.45e-9$ | $9.31e-9$ |
| 50 | 28 | 0.664 | 0.501 | 0.143 | $3.73e-9$ | $7.45e-9$ |
| 52 | 28 | 0.701 | 0.451 | 0.236 | $3.73e-9$ | $7.45e-9$ |
| 56 | 28 | 0.849 | 0.648 | 0.187 | $3.73e-9$ | $7.45e-9$ |

As a secondary measure of state-specific representational strength, we quantified a template-specific contribution to global explained variance (GEV) as follows:

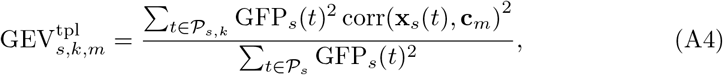

where *𝒫*_*s*_ denotes all GFP peaks for subject *s, 𝒫*_*s,k*_ the GFP peaks assigned to local minimum *k*, and **c**_*m*_ the *m*-th canonical microstate template. This metric captures the GFP-weighted fit of template *m* to the peak maps assigned to state *k*, expressed relative to the subject’s total GFP-peak variance. Accordingly, the values reflect both template fit and the overall contribution of that state to subject-level variance, and are not intended to be interpreted as the total explained variance of a canonical microstate class. Template-specific GEV contributions were broadly consistent with the correlation-based ranking (Fig. A2b): the *D*-like family (7, 56) showed the largest contributions, the *B*-like families (13, 50) and (11, 52) showed intermediate contributions, and the weak family (26, 37) remained uniformly low. This convergence across complementary metrics supports the topographic assignments derived from the spatial-correlation analysis.

## Footnotes

1 https://github.com/ThomasKoenigBern/MS-Template-Explorer

